# A new triple fluorescence reporter system for discrimination of Apobec1 and Apobec3 C-to-U RNA editing activities and editing-dependent protein expression

**DOI:** 10.1101/2021.03.03.433736

**Authors:** Barbara Schweissthal, Kea Brunken, Julia Brach, Leonie Emde, Florian Hetsch, Steffen Fricke, Jochen C. Meier

## Abstract

The human body is composed of many different cell types which communicate with each other. In particular, the brain consists of billions of neurons and non-neuronal cells which are interconnected and require tight and precise regulation of cellular processes. RNA editing is a cellular process that diversifies gene function by enzymatic deamination of cytidine or adenine. This can result in changes of protein structure and function. Altered RNA editing is becoming increasingly associated with all kind of disease, but most approaches use advanced sequencing technologies to analyze bulk material. However, it is also becoming progressively evident that changes in RNA editing have to be analyzed and considered in a cell type specific way. We present here a triple fluorescence reporter system that discriminates between Apobec1- and Apobec3-dependent C-to-U RNA editing at the single cell level. In particular, the Apobec3 reporter enables C-to-U RNA editing inducible protein expression through generation of a RNA splice donor site. We used the new system here to analyze Apobec1- and Apobec3-dependent RNA editing in primary neuron culture. The results reveal a large heterogeneity of C-to-U RNA editing in neurons and glia cells, and they show that GABAergic neurons are not able to perform Apobec1-dependent RNA editing, but Apobec3-dependent editing. Altogether, the new system can be the foundation of therapeutic application systems that counteract changes in Apobec3-dependent RNA editing in disease while simultaneously monitoring Apobec1-dependent RNA editing at the single cell level.

## Introduction

RNA editing diversifies genome-coded protein functions by enzymatic deamination of cytidine or adenine nucleobases in mRNA. In recent years, it turned out that alterations in RNA editing may play a key role in a plethora of idiopathic human diseases [for overview see (Meier et al., 2016; Krestel and Meier, 2018)]. In particular, recent experimental evidence suggests that increased mRNA editing in the hippocampus is involved in epilepsy (Eichler et al., 2008; Srivastava et al., 2017). However, experiments also demonstrated that functional consequences of changes in RNA editing have to be investigated at the single cell level because cell type-specific expression of a C-to-U RNA edited target in a transgenic mouse model of epilepsy revealed striking differences in the symptomatology of the mice (Winkelmann et al., 2014; Caliskan et al., 2016). Thus, molecular tools for detection of RNA editing at the single cell level are required. Some tools for monitoring C-to-U RNA editing at the single cell level have already been presented (Kankowski et al., 2018; Severi and Conticello, 2015). More recently, a new molecular tool for C-to-U RNA editing dependent protein expression was presented (St.Martin et al., 2018), but in this case some additional artificial sequences remain after C-to-U RNA editing and subsequent Cas9-dependent processing of the mRNA.

We present here a novel molecular tool for C-to-U RNA editing dependent protein expression that does not involve additional artificial sequences expressed in the protein of interest if RNA editing occurs. The tool consists of two fluorescent proteins separated by a mutated Apobec1 consensus sequence directly followed by a synthetic intron. C-to-U RNA editing of the consensus sequence generates a RNA splice donor site that is used in combination with a constitutive downstream splice acceptor site. This construct mediates C-to-U RNA editing induced mRNA splicing and resulting protein expression due to removal of STOP codons in the synthetic intron. However, modification of the Apobec1 consensus sequence required for generation of the editing induced splice donor site makes this construct accessible for Apobec3 dependent RNA editing. In conjunction with our Apobec1 specific RNA editing sensor tool (Kankowski et al., 2018) in a single DNA plasmid separated by an IRES our new triple fluorescence reporter with Apobec3 RNA editing induced protein expression can distinguish Apobec1 and Apobec3 mediated C-to-U RNA editing at the single cell level. Using this new tool, the results show that there is great heterogeneity between Apobec1 and Apobec3 dependent RNA editing in primary hippocampal glia cells and neurons. In particular, we also show that GABAergic neurons selectively perform Apobec3 dependent RNA editing. Altogether, our system enables Apobec3-dependent protein expression and monitoring of Apobec1-and Apobec3-dependent C-to-U RNA editing at the single cell level. These results are the foundation of therapeutic application systems that counteract changes in Apobec3 dependent RNA editing in disease while simultaneously monitoring Apobec1 dependent RNA editing.

## Materials and Methods

### Molecular biology

The Apobec1-specific RNA editing sensor construct is published (Kankowski et al., 2018). Based on this published construct, generation of the Apobec3 dependent protein expression system was established. For this purpose, a synthetic construct was synthesized by Biomatik (Cambridge, Ontario, N3H 4R7, Canada) and delivered subcloned in a pBluescript vector. The construct contains a core sequence with a synthetic intron directly downstream of the Apobec1 consensus (Kankowski et al., 2018). The intronic core sequence is composed of a short intronic GlyR β intron (Winkelmann et al., 2015) and a modified gephyrin intronic sequence (Förstera et al., 2010). Based on this construct we modified the Apobec1 consensus sequence by introducing mutations T→G at −2 and +3 positions relative to the C-to-U RNA edited site. We did this because maximum entropy model (Yeo and Burge, 2004) predicted a strong splice donor site if C-to-U editing occurs and if a guanine is present at position −1 (Table 1). Of course, constructs with an adenine instead of guanine at position −1 relative to the edited position served as negative controls for editing dependent generation of the splice donor site. To enable C-to-U RNA editing dependent protein expression (here mCherry), a cassette coding for 2A peptide (Tang et al., 2009) and downstream a mCherry-coding open reading frame was inserted at the multiple cloning site at the 3’ end of the synthesized construct using standard restriction enzyme dependent molecular cloning. Furthermore, for some experiments, the GlyR α3 TM3-4 intracellular loop was deleted using fusion-PCR cloning. Finally, we generated the triple fluorescence reporter with the Apobec1-specific RNA editing sensor tool (translocation of eGFP into the nucleus indicates RNA editing) and the Apobec3 C-to-U RNA editing inducible protein expression system (constitutive miRFP703 expression and editing induced mCherry expression) separated by an IRES using NEBuilder® HiFi DNA Assembly technology. Plasmid pmiRFP703-N1 was a gift from Dr. Vladislav Verkhusha (Addgene plasmid # 79988; http://n2t.net/addgene:79988; RRID:Addgene_79988) (Shcherbakova et al., 2016).

**Table 1:** Prediction of splice sites according to maximum entropy model with numbering of splice sites according to Figure 2.

| Splice sites | Mutations | Maximum Entropy Model to score |
| --- | --- | --- |
| splice site 1 (donor) | / | +8.41 |
| splice site 2 (donor) | GC | +1.96 |
|  | GC, -2 T→G | +8.88 |
|  | GC, +3 T→G | +9.11 |
|  | GC, -2/+3 T→G | +11.08 |
| splice site 3 (acceptor) | / | +11.50 |
| splice site 4 (acceptor) | / | +4.62 |

For assessment of C-to-U RNA editing dependent splicing and resulting protein (mCherry) expression total RNA was isolated from HEK293T and HepG2 cells using TRIzol reagent (Invitrogen). To eliminate residual plasmid DNA, the sample was treated with RNase free DNase (catalog no. 11119915001, Roche) as described previously (Knoll et al., 2019). Briefly, 10 units were incubated (20 min, 37 °C) with 50 μg RNA sample and purified using RNeasy Protect Mini Kit (catalog no. 74124, Qiagen) according to manufacturer’s instructions. Complementary DNA was obtained by reverse transcription (Superscript II, catalog no. 18064014, Invitrogen) of 2 μg RNA with an equimolar mixture of 3’-anchored poly-T oligonucleotides (T18V, T15V, T13V). DNA-free isolation was verified by PCR amplification (RedTaq® Polymerase, catalog no. D4309, Sigma-Aldrich). For this purpose, different combinations of oligonucleotides covering non-transcribed and transcribed regions of the plasmids were used, as specified in figure legends. To assess whether Apobec3 edits transfected plasmid DNA we isolated the transfected plasmids using Invisorb® Spin Plasmid Mini Two (INVITEK Molecular GmbH, Germany) including RNAse digest during plasmid isolation. PCR was performed to amplify regions of interest followed by DNA sequencing using mix2seq kit (Eurofins).

### Cell lines, primary neuron cultures, microscopy and morphometry

Transient protein expression of constructs in HEK293T, HepG2 cell lines, and primary hippocampal cell cultures was performed as described previously (Kankowski et al., 2018), except that mouse hippocampus was used. In brief, primary hippocampal neurons were prepared from NMRI mice at embryonic day 18 and plated on NaOH treated and poly-L-lysine coated coverslips at a density of 5×10^4^ cells per well. Following one (cell lines) or 3 days expression (primary hippocampal cultures) cells were fixed using paraformaldehyde and coverslips mounted using Vectashield (Vector Laboratories, Burlingame, CA, USA) with or without DAPI (cell lines). Primary hippocampal cultures were further processed for immunochemistry using antibodies (anti-GABA 1:750, Sigma, with anti-rabbit IgG AMCA 1:200, Dianova, and anti-MAP2 1:500, Synaptic Systems, with anti-guinea pig IgG AMCA 1:200, Dianova) and mounted using Vectashield without DAPI.

Fluorescence microscopy was performed using either Olympus BX51 upright microscope (metal halide illumination) or Nikon ECLIPSE Ni-U upright microscope (pE-4000 fluorescence LED system) for acquisition of miRFP703, eGFP, and mCherry signals. Using LED illumination, the excitation wavelengths were trimmed to 660 nm (miRFP703), 595 nm (mCherry), and 470 nm (eGFP). GABA and MAP2 signals were acquired using 395 nm excitation wavelength. For each experiment, illumination strength and exposure time were adjusted and kept constant to avoid saturated signal intensities and to be able to unambiguously be able to compare fluorescence intensity ratios. Using software Metamorph (Molecular Devices, Sunnyvale CA) or NIS Elements (Nikon GmbH Microscope Solutions, Düsseldorf, Germany), line scans were applied so that the lines covered cytosol and/or nucleus of transfected cells. Fluorescence intensity values along the scanned lines were exported to Excel software (Microsoft, Redmond WA, USA), and the pixel intensity values were separated into cytosolic and nuclear parts of the cell according to the DAPI signal (Apobec1 editing sensor) or nuclear regions only (Apobec3 inducible mCherry expression).

## Results

We modified the recently described Apobec1 editing sensor domain (Kankowski et al., 2018) at different positions relative to the edited cytidine (−1 A → G (“AC” versus “GC”) and −2/+3 T → G). Compared to the previously published sensor domain (Kankowski et al., 2018), the APOBEC1 consensus sequence was thus differentially mutated at three positions and is followed downstream by a synthetic intron derived from *Glrb* and *Gphn* introns (Förstera et al., 2010; Winkelmann et al., 2015), see Figures 1 and 2. Mutation at position −1 is obligatory if C-to-U RNA editing is supposed to generate a splice donor site (GU). Mutations T → G at sites −2 and +3 were predicted (Table 1) using maximum entropy model (Yeo and Burge, 2004) and hence were mutated accordingly. The eight different constructs were expressed in HEK293T cells and RNA was isolated one day following transfection. RNA was treated with DNase to eliminate residual plasmid DNA in RNA preparations. cDNA was produced, amplified with PCR (Figure 1) and sequenced. Amplification of plasmid DNAs was performed in order to verify functionality of oligonucleotides and elimination of plasmid DNA in the RNA preparations. Four results are intriguing: i) In contrast to the predicted splice acceptor site at the end of the synthetic intron a splice site at the end of the transmembrane domain of the GlyR α3k-loop was used by the splice machinery, ii) only the combination of −2/+3 T → G enabled C-to-U RNA editing-dependent RNA splicing in constructs with G at position −1 (for better overview see Figure 2), iii) Apobec1 80M or 80I variants do not seem to have a different impact on editing-induced RNA splicing, and iv) C-to-U RNA editing did not occur when constructs with adenine at −1 were expressed; furthermore, editing required −2/+3 T → G substitutions.

**Figure 1:**
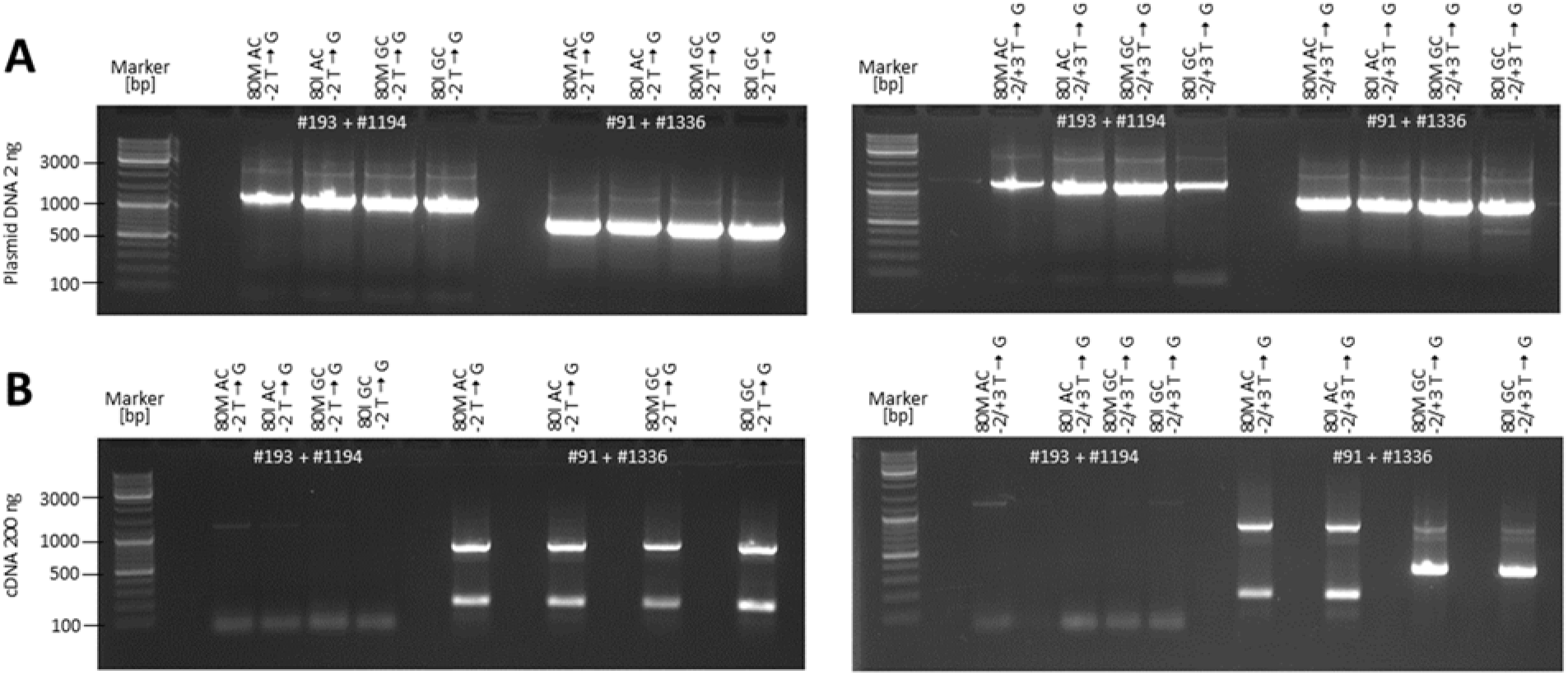
PCR amplification of plasmid DNA **(A)** used for transfection of HEK293T cells or of cDNA **(B)** of mRNA isolated out of the transfected HEK293T cells. **(A)** Amplification of plasmid DNAs. Oligonucleotide #193 (5’-GGAGGTCTATATAAGCAGAGC-3‘) binds in the CMV promotor region and was used in combination with #1194 (5’-ATGGCTCTGTTGCTGAGATGTC-3‘) binding at the N-terminus of Apobec1. Oligonucleotide #91 (5’-CATGGTCCTGCTGGAGTTCGTG-3‘) binds at the C-terminus of eGFP and was used in combination with #1336 (5’-GCTGATGAATGTCTTCATGCC-3‘) binding in between the TM4 of the α3k-loop and the 2A-peptide. Thus, PCR using #193 + #1194 will only amplify DNA products if DNA is present, while the combination #91 + #1336 amplifies cDNA. **(B)** Amplification of cDNA from transfected HEK293T cells. Note that #193 + #1194 yields faint PCR bands, indicating that the cDNA was almost completely free of plasmid DNA. On the other hand, #91 + #1336 yields prominent PCR bands with different sizes that were excised, purified and DNA sequenced. Note that mRNA of transfected clones coding for 80M or 80I Apobec1 and mutated at −1 A → G (“GC”) and −2/+3 T → G were differently processed than all other mRNAs, indicating that C-to-U RNA editing-dependent RNA splicing occurred; see Figure 2 for an overview of splice sites that were used.

**Figure 2:**
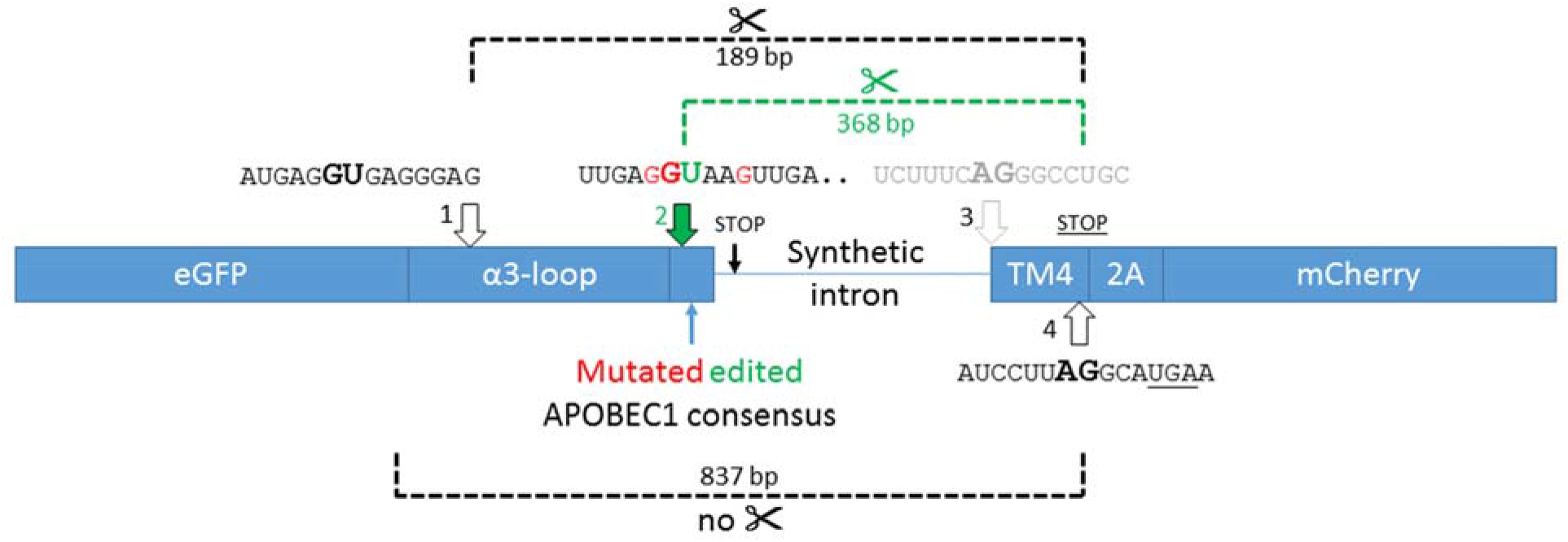
Scheme depicting the working principle of C-to-U RNA editing inducible protein expression (CUREIPES). The editing sensor domain is located downstream of eGFP. Compared to the previously published sensor domain (Kankowski et al., 2018), the APOBEC1 consensus sequence was mutated at three positions (red) and is followed downstream by a synthetic intron derived from *Glrb* and *Gphn* introns (Winkelmann et al., 2015; Förstera et al., 2010). If C-to-U RNA editing occurs, the splice site (2) **GU** is created and used in combination with splice site (4), leading to elimination of the first STOP codon and resulting in an open reading frame with 2A_mCherry located downstream of the TM4 domain of the GlyR α3k loop. Due to frame shift, the second <u>STOP</u> codon is no longer a STOP codon. Splicing at the predicted site (3) does not occur. If there is no C-to-U RNA editing, splice sites (1) and (4) are used (✂), leading to frame shift and protein truncation (<u>STOP</u>) immediately downstream of splice site (4). If there is no splicing (no ✂), the STOP codon at the begin of the synthetic intron leads to protein truncation. Dashed brackets indicate the length of PCR products obtained from DNA-free RNA preparations of transfected HEK293 cells that were verified by DNA sequencing.

To verify C-to-U RNA editing-dependent RNA splicing and resulting protein (mCherry) expression from the new system we termed CUREIPES, we performed fluorescence microscopy using Apobec1 80I coding constructs (Figure 3A, B). Furthermore, the GlyR α3k-loop was deleted in order to assess whether it plays a role in CUREIPES (Figure 3C, D).

**Figure 3:**
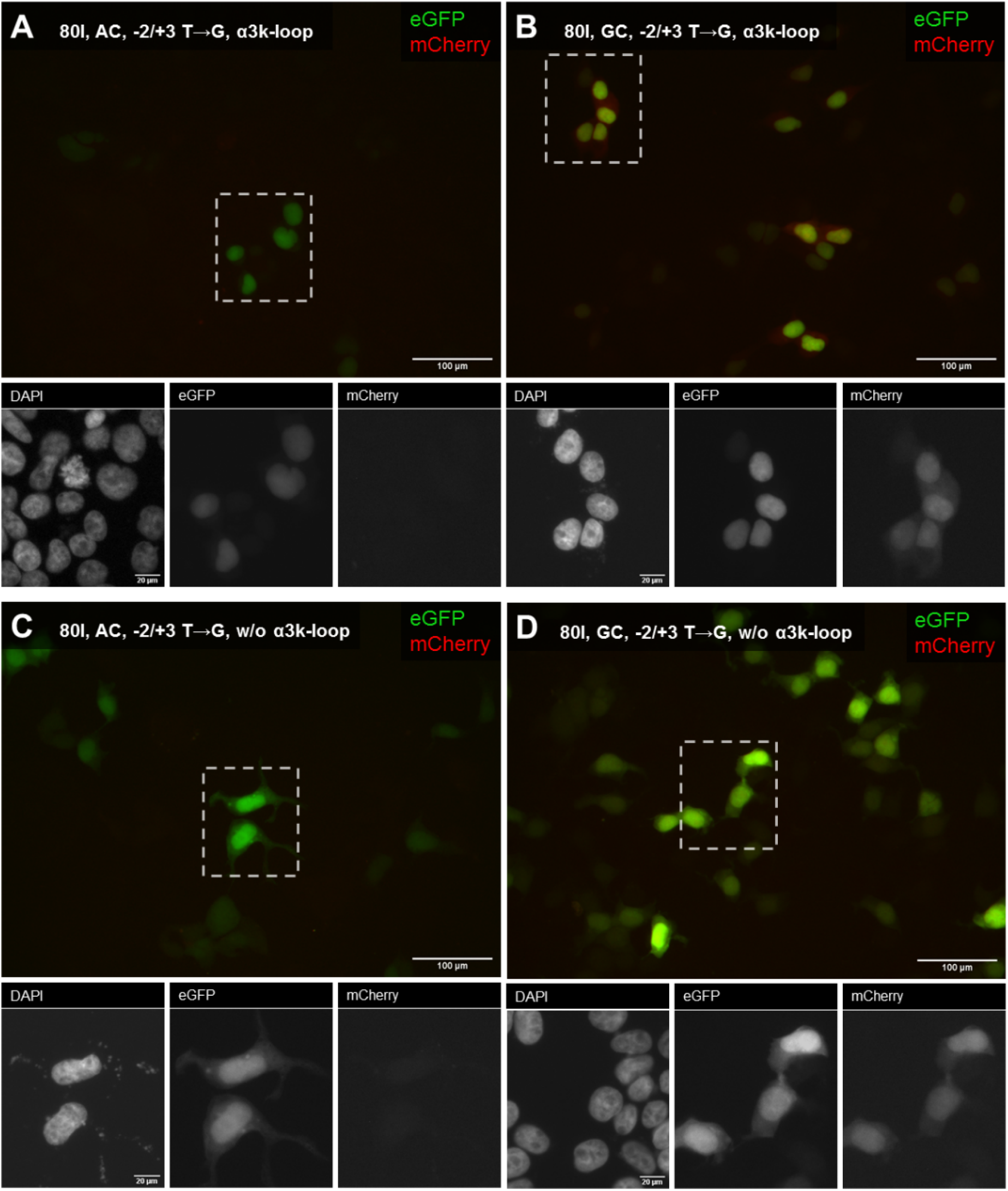
Fluorescence microscopy of transfected HEK293T cells using the indicated constructs with **(A, B)** GlyR α3k-loop or without (“w/o”) the loop **(C, D)**. Clearly, quantification of the ratio of fluorescence intensities between mCherry and eGFP indicate that mCherry was expressed if constructs −2/+3 T → G with G at position −1 were expressed (“GC”). In contrast, “AC” constructs with A at position −1 completely abolished C-to-U RNA editing-induced RNA splicing and prevented mCherry protein expression.

We next wondered whether CUREIPES functions in HEK293T cells if Apobec1 and ACF are not co-expressed via 2A-peptides. For this purpose, we generated constructs −2/+3 T → G with G at position −1 with or without GlyR α3k-loop and deleted ACF/Apobec1 (Figure 4).

**Figure 4:**
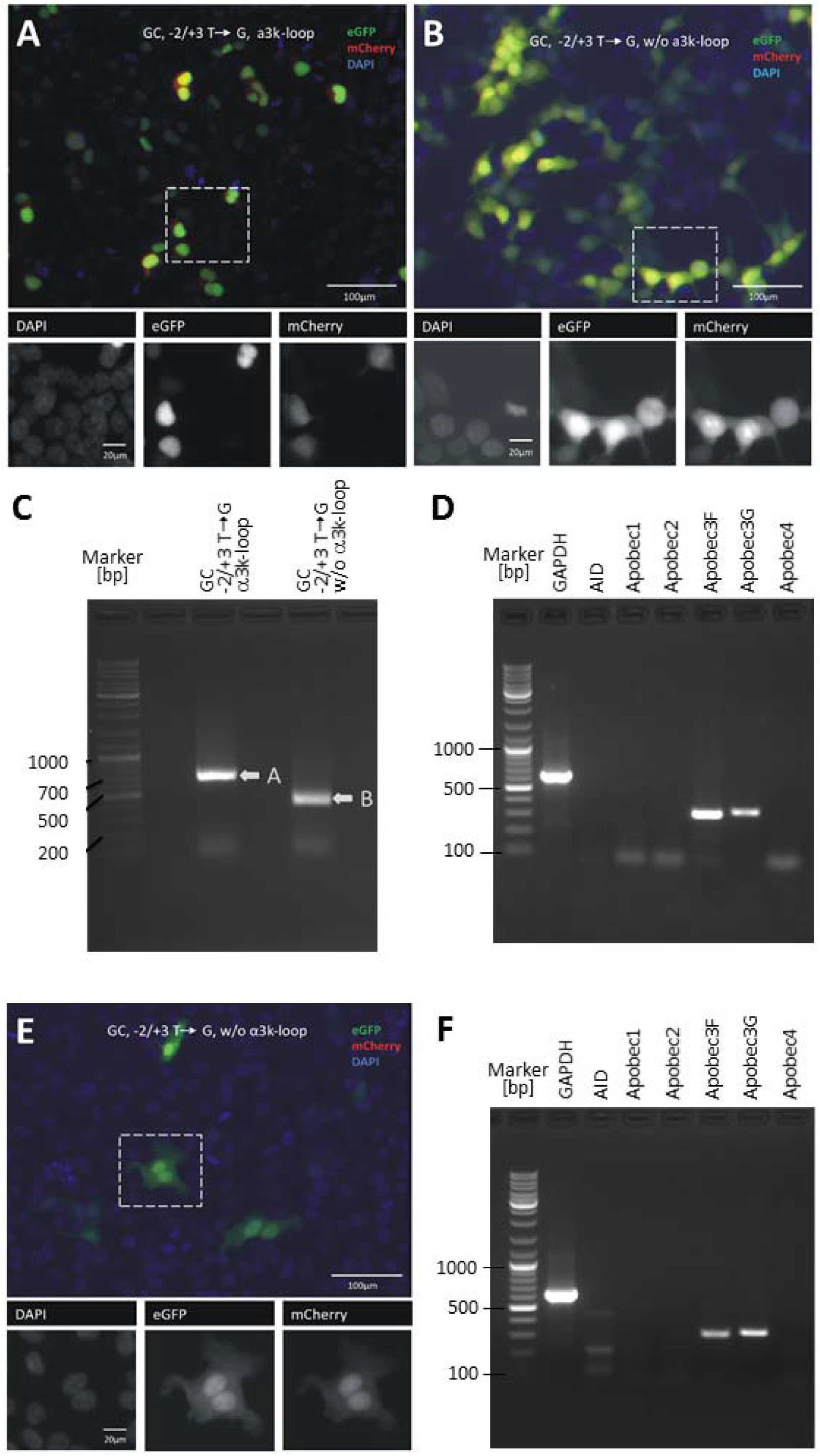
Fluorescence microscopy and molecular profiling of transfected HEK293T and HepG2 cells. **(A-D)** HEK293T cells. Constructs with GlyR α3k-loop **(A)** or without (“w/o”) the loop **(B)** were expressed in HEK293T cells. Clearly, mCherry expression is observed irrespectively of the GlyR α3k-loop. **(C)** We analyzed again cDNA from transfected HEK293T cells and excised PCR bands (band “A” and “B”) obtained with PCR amplification using oligonucleotides binding in the middle of eGFP open reading frame 5’-GGCATCGACTTCAAGGAGG-3’ and binding at the junction between 2A-peptide and mCherry open reading frame 5’-CGACCGGTGGATCCAGAGC-3’. DNA sequencing confirmed C-to-U RNA editing-dependent RNA splicing. **(D)** Profiling of HEK293T cells with regard to cell-endogenous expression of different C-to-U RNA editing enzymes. **(E, F)** HepG2 cells. The construct without (“w/o”) the GlyR α3k-loop was used **(E)**. **(F)** Profiling of HepG2 cells with regard to cell-endogenous expression of different C-to-U RNA editing enzymes. Oligonucleotides used for molecular profiling of HEK293T and HepG2 cells were as follows: housekeeping gene product GAPDH (5’-ATGGCACCGTCAAGGCTGAG-3’ and 5’-CGACGCCTGCTTCACCACC-3’), activation-induced cytidine deaminase AID (5’-GTCCGCTGGGCTAAGGGTC-3’ and 5’-GCACAGTCGTAGCAGGGGC-3’), Apobec1 (5’-CTTCAACCGGTGACCCCACTC-3’ and 5’-TGCGTACAACATCATCCACAGAGG-3’), Apobec2 (5’-TGATTGAGCTGCCGCCCTTTG-3’ and 5’-CAGGTTCTTGGTCTTGCTAAGGG-3’), Apobec3F (5’-GCCTCACTTCAGAAACACAGTGG-3’ and 5’-CAGCTTCGCCACACAGTCCG-3’), Apobec3G (5’-GCCTCACTTCAGAAACACAGTGG-3’ and 5’-AACGTGGCCATATCCCTTGTAC-3’), and Apobec4 (5’-TAGCCACACCTCAGCAAGGG-3’ and 5’-TGCAACTGCTAGCATGGCCC-3’).

HEK293T cells obviously edit mRNAs coding for CUREIPES irrespectively of the presence of the GlyR α3k-loop (Figure 4A, B), and sequencing of PCR-amplified CUREIPES-coding cDNA (Figure 4C) proved it at the molecular level. This result is surprising because HEK293T cells are not supposed to express Apobec1. Furthermore, we wanted to know whether CUREIPES would also function in the human hepatoma cell line HepG2 (Figure 4E). HepG2 cells were widely used to study Apobec1-dependent C-to-U RNA editing, and this cell line does not express Apobec1 (Chen et al., 1987; Teng et al., 1993; Chen et al., 2010). The results in Figure 4E demonstrate that CUREIPES is edited also in HepG2 cells. Again, sequencing of PCR-amplified CUREIPES-coding cDNA proved it at the molecular level.

We characterized the expression pattern of AID/Apobec variants in CUREIPES-expressing cells treated with 1-(β-D-Ribofuranosyl)-2(1H)-pyrimidinone (zebularine, Sigma-Aldrich CAS 3690-10-6, final concentration 50 μM in 0.1% DMSO), a nucleoside analog of cytidine that binds to the active site of cytidine deaminases (Vincenzetti et al., 2000) and was proposed to act as transition state analog inhibitor (Zhou et al., 2002) of these enzymes. Alternatively, zebularine can also act as DNA methylation inhibitor. For control purpose, we treated cells with 0.1% DMSO in the absence of zebularine. Again, sequencing of amplified CUREIPES revealed C-to-U RNA editing but ruled out an effect of zebularine on C-to-U RNA editing of CUREIPES in both HEK293T and HepG2 cells. Molecular profiling of the expression of C-to-U RNA editing enzymes in transfected cells (HEK293T and HepG2) in the presence or absence of zebularine showed that these cells only expressed Apobec3 (Figure 4D, F, respectively). Several reports suggested that HEK293T cells do not endogenously express Apobec3 because antiviral Apobec3 effects required overexpression of that protein [e.g.(Sheehy et al., 2002; Shindo et al., 2003)]. It is possible that our HEK293T cell clone is particular in this respect. Therefore, Apobec3 expression was also assessed in an independent HEK293T cell clone at the University of Mainz, Germany. However, this additional control experiment confirmed our results (data not shown). Furthermore for control purpose, we assessed HEK293T cell-endogenous expression of Apobec3 in non-transfected cell as well as in cells expressing CUREIPES with or without the GlyR α3-loop (data not shown). This experiment ruled out the possibility that transfection of the HEK293T cells per se and expression of CUREIPES in particular triggered HEK293T cell-endogenous Apobec3 expression. Clearly, HEK293T cells expressed Apobec3, and in particular 3B, 3C, 3D1/2, 3F1/2, 3G1, and 3H, while HepG2 cells expressed Apobec3A, 3B, 3C, 3F1/2, 3G1, and 3H (Suppl. Figure 1). Thus, Apobec3 variants expressed in the transfected HEK293T and HepG2 cells are Apobec3B, 3C, 3F1, 3F2, 3G1, and 3H. As Apobec3F and G variants were suggested to preferentially edit DNA but can also edit mRNA (Apobec3G) [(Siriwardena et al., 2016) and references therein; (Sharma et al., 2016)], we isolated the transfected plasmid DNAs out of the transfected HEK293T cells using a plasmid mini-preparation kit for plasmid DNA isolation out of bacteria in the presence of RNase. PCR amplification of the purified plasmids was performed using oligonucleotide (5’-GGAGGTCTATATAAGCAGAGC-3‘) binding in the CMV promotor and reverse oligonucleotide 5’-CGACCGGTGGATCCAGAGC-3’ binding at the junction between 2A-peptide and mCherry open reading frame. Sequencing of the amplified plasmid DNA using oligonucleotide 5’-GGCATCGACTTCAAGGAGG-3’ binding in the middle of eGFP open reading frame ruled out C-to-T double-stranded DNA editing by HEK293T cell-endogenous Apobec3. We are currently performing shRNA-mediated knockdown experiments in HEK293T cells to determine which Apobec3 variant is responsible for editing-dependent RNA splicing in CUREIPES.

Taken together, the results obtained so far suggest that the Apobec1 consensus sequence modified at positions −2 and +3 (T→G) and −1 (A→G) is used for C-to-U RNA editing-dependent RNA splicing and results in protein (mCherry) expression in HEK293T which endogenously express different Apobec3 variants. To explore and eventually discriminate Apobec1-and Apobec3-dependent C-to-U RNA editing we substituted miRFP703 for eGFP in the CUREIPES construct and generated a new vector with CUREIPES and the Apobec1-specific C-to-U RNA editing sensor (Kankowski et al., 2018) separated by an IRES (Figure 5A). The resulting construct encodes a triple fluorescence sensor for detection of Apobec1-dependent C-to-U RNA editing (indicated by translocation of eGFP from cytosol to nucleus) and Apobec3-dependent mCherry expression (miRFP703 is constitutively expressed). This construct was first expressed in HEK293T cells (Figure 5B, C). Altogether, Figure 5 indicates Apobec3-dependent RNA editing in agreement with the results described above. Regarding the Apobec1-specific RNA editing sensor (eGFP nucleus/cytosol), the results in Figure 5 are also consistent with previously reported data (Kankowski et al., 2018). Thus, the triple fluorescence reporter is able to discriminate Apobec1- and Apobec3-dependent C-to-U RNA editing in HEK293T cells.

**Figure 5:**
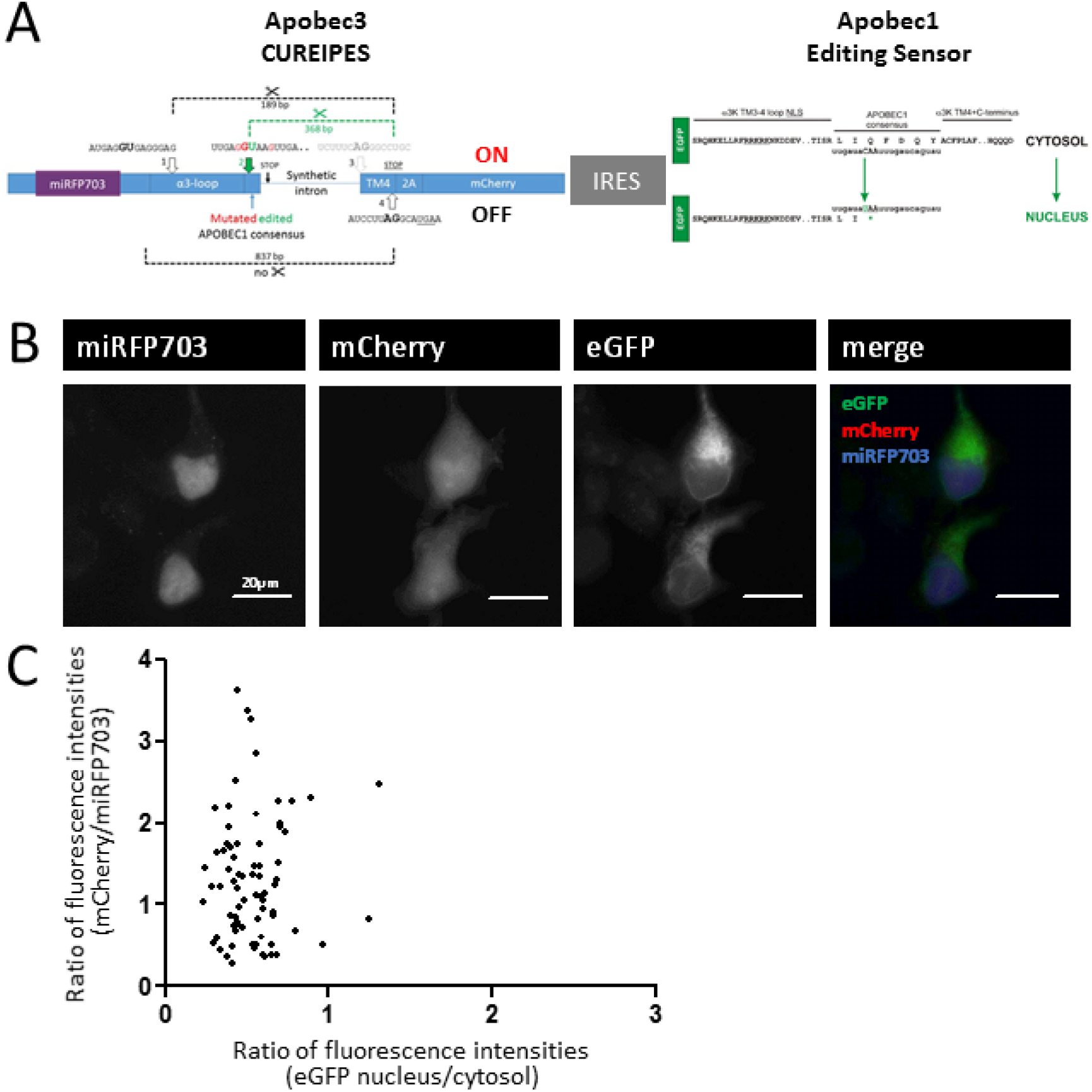
**(A)** Scheme depicting design and work principle of the triple fluorescence construct (note that Apobec1 Editing Sensor was published previously; Kankowski et al., 2008). **(B)** Fluorescence microscopy of transfected HEK293T cells expressing the triple fluorescence reporter. **(C)** Scatter blot of fluorescence ratios indicating Apobec1-(eGFP nucleus/cytosol) and Apobec3-dependent (mCherry/miRFP703) C-to-U RNA editing. Each dot represents a single cell.

In contrast to HEK293T cells, primary hippocampal cultures composed of different cell types express AID, Apobec1, 2, 3, and 4, as was determined using bulk RNA material (Kankowski et al., 2018). Hence, we expressed the triple fluorescence reporter in mouse primary hippocampal cultures at day in vitro 7 (DIV7) and identified neurons using MAP2-staining, GABAergic neurons using GABA-staining, and MAP2-negative glia cells 3 days later in order to provide information about Apobec1- and Apobec3-dependent RNA editing at the single cell level in immunochemically identified different cell types (Figure 6).

**Figure 6:**
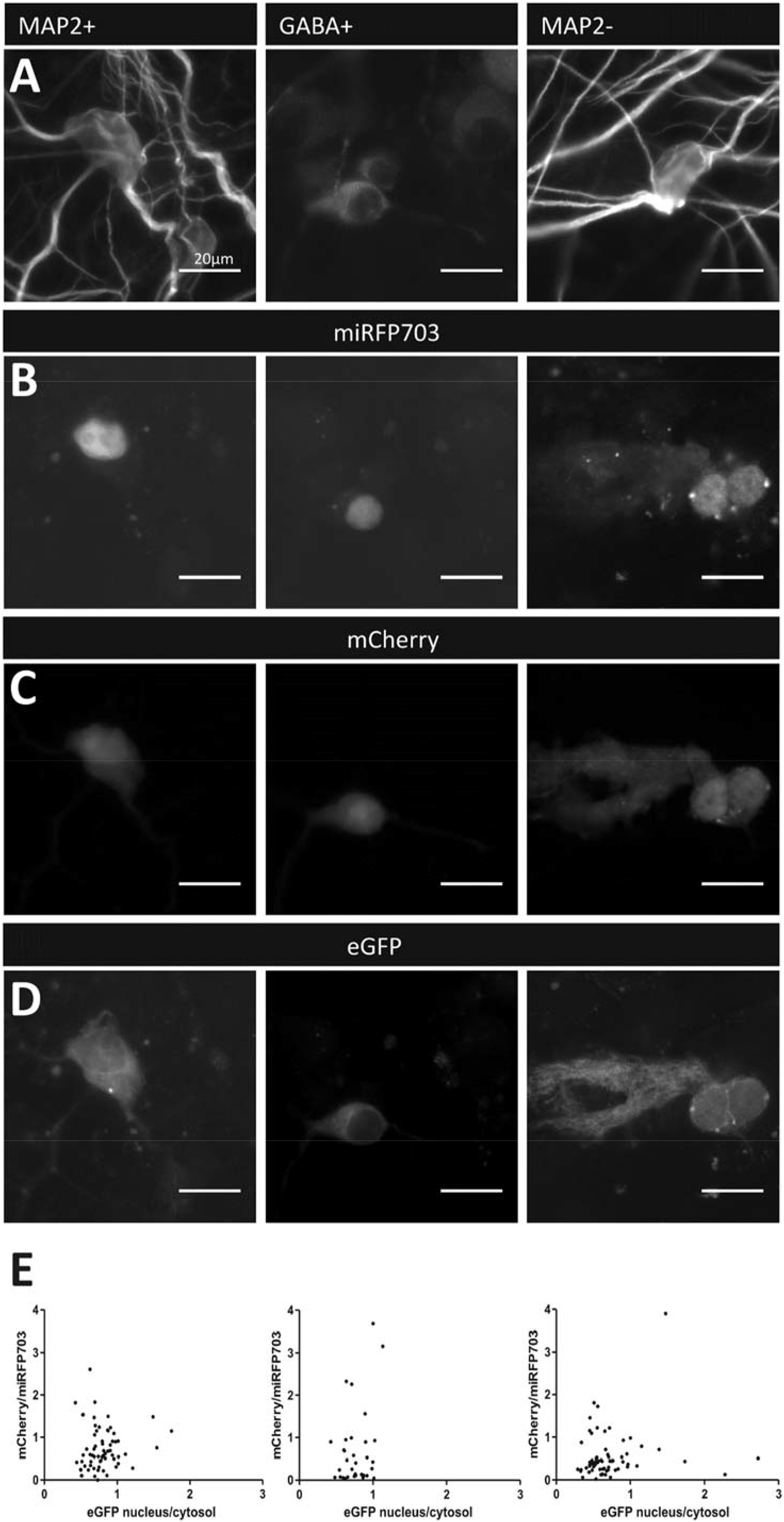
Fluorescence microscopy of transfected mouse primary hippocampal cultures expressing the triple fluorescence reporter. **(A)** Representative images showing MAP2-positive (“MAP2+”), GABA-positive (“GABA+”), and MAP2-negative (“MAP2-“) cell types in the transfected cultures. **(B-D)** Representative images of showing miRFP703 **(B)**, mCherry **(C)**, and eGFP **(D)** signals **(E)** Scatter blot of fluorescence ratios representing Apobec1- and Apobec3-dependent C-to-U RNA editing (mCherry/miRFP703 and eGFP nucleus/cytosol, respectively). Each dot represents a single cell.

Results shown in Figure 6 reveal cell type-specific heterogeneity of Apobec1- and Apobec3-dependent C-to-U RNA editing. In particular, GABAergic neurons do not show Apobec1-dependent RNA editing, while some cells of this cell type show CUREIPES signals. In contrast, neurons and glia cells display a large heterogeneity regarding their individual signals corresponding to Apobec1- and Apobec3-dependent C-to-U RNA editing.

## Discussion

The results in this original research paper presents a new molecular tool for C-to-U RNA editing-dependent protein expression. In combination with the recently published Apobec1-specific RNA editing sensor (Kankowski et al., 2018), the C-to-U RNA editing-inducible protein expression system CUREIPES is able to discriminate between Apobec1- and Apobec3-dependent RNA editing.

Based on the Apobec1-specific consensus sequence (Kankowski et al., 2018) we introduced mutations at positions −2 and +3 relative to the edited cytidine and furthermore substituted guanine for adenine at position −1 to allow editing-dependent generation of a splice donor site (Table 1, No 2 in Figure 2). While mutations around the edited cytidine are in agreement with prediction of splice sites with regard to usage of the editing-dependent splice donor site, predictions regarding the splice acceptor site are not in agreement with our actual results (Table 1). Instead of the predicted splice acceptor site (No 3 in Figure 2) at the end of the intronic core sequence composed of a short intronic GlyR β intron (Winkelmann et al., 2015) and a modified gephyrin intronic sequence (Förstera et al., 2010), a splice acceptor site within TM4 of GlyR α3 is used (No 4 in Figure 2). It is intriguing to see that an adenine at position −1 completely abolished C-to-U RNA editing, and that −2/+3 T → G substitutions were necessary for Apobec3-dependent C-to-U RNA editing. Furthermore, if C-to-U RNA editing-dependent creation of the splice donor site is not possible (in AC constructs), a non-predicted splice donor site at the beginning of the GlyR α3k-TM3/4-loop is used. These results are surprising and point to the possibility that GlyR α3k full-length protein expression in neuronal tissue is subject to unpredicted RNA splicing and hence, expression of truncated GlyR α3 proteins.

The new triple fluorescence reporter indicates Apobec1-specific RNA editing and enables Apobec3-dependent protein expression at the single cell level and in a cell type-specific way. We can rule out that Apobec1 edits CUREIPES because if this was the case every cell in the primary mouse hippocampal cultures that shows Apobec3-dependent RNA editing should also show Apobec1-dependent translocation of the sensor into the nucleus – this was not the case. The results in our study clearly add a valuable molecular tool for the study of Apobec1- and Apobec3-dependent C-to-U RNA editing at the single cell level, which can nicely complement and enhance research approaches that use tissue bulk material for advanced sequencing analyses in health and disease [e.g. (Srivastava et al., 2017)]. Actually, the results presented here showed that GABAergic neurons play a particular role as no Apobec1-dependent but Apobec3-dependent C-to-U RNA editing was detected in this cell type. MAP2-positive and −negative cell types furthermore displayed a great heterogeneity with regard to Apobec1/3-dependent RNA editing. Thus, cell type-specific approaches are clearly required to address the role of RNA editing specifically at the single cell level – our previous research indeed demonstrated neuron type-specific roles of increased C-to-U RNA editing of a single gene product in the context of mesial temporal lobe epilepsy (Eichler et al., 2008; Winkelmann et al., 2014; Caliskan et al., 2016). Now, we have to identify more molecular targets of aberrant C-to-U RNA editing and enzymes involved using cell type-specific approaches to identify key molecular players in development and disease. In particular, the lack of model systems for the study of Apobec3-dependent editing in a cross-species-valid way represented a major burden for the study of Apobec3-dependent mechanisms (Siriwardena et al., 2016). As the consensus site in CUREIPES is not compatible with consensus site sequence analyses of Apobec3G (Sharma et al., 2016; Rausch et al., 2009) in DNA templates our results suggest that, in human cells, CUREIPES mRNA is edited by Apobec3B, 3C, 3F, or 3H rather than by Apobec3G in HEK293T and HepG2 cells. This is compatible with conserved RNA binding motifs present in these Apobec3 variants (Zhen et al., 2012). A major advantage of our new system over other systems [recently reviewed in (Chieca et al., 2021)] is that there is almost no additional protein sequence (except PVD; P: very end of 2A-peptide, and VD: *Sal*I restriction enzyme site) that will be expressed in response to Apobec3 RNA editing-dependent expression of selected proteins of interest. Taken together, CUREIPES is functional in rodent and human cellular systems and hence, provides the first molecular tool for the study Apobec3-dependent mechanisms at the single cell level and across species.

CUREIPES may open avenues for future research including developmental profiling of C-to-U RNA editing in any tissue or specifically in particular cell types or brain regions if the CMV promotor is substituted. Furthermore, it will be possible now to counteract disease-associated changes in RNA editing by substituting any therapeutically relevant protein (including RNA editing regulatory proteins) for mCherry. It is also conceivable to combine established inducible protein expression systems (e.g. TetON/OFF) with CUREIPES by substituting Tet-regulatory proteins for mCherry. Other possible applications include C-to-U RNA editing-dependent gene regulation (transcription factors in place of mCherry), gene editing (e.g. CRISPR), or cell type specific C-to-U RNA editing-dependent generation of knockout animals (e.g. Cre recombinase). Indeed, any application reviewed recently for the case of A-to-I RNA editing-dependent systems (Aquino-Jarquin, 2020) could now also be applied using our C-to-U RNA editing-specific CUREIPES.

Taken together, this study presents a new triple fluorescence reporter for monitoring Apobec1- and Apobec3-dependent C-to-U RNA editing at the single cell level in rodent and human cells. Furthermore, CUREIPES is a new system that enables Apobec3 C-to-U RNA editing-dependent expression of any protein of interest, which should be useful for a very wide range of basic research and therapeutic applications.

## Acknowledgements

We thank Vanessa Hering (Technical University Braunschweig) for excellent technical assistance. We thank Dr. Vladislav Verkhusha (Albert Einstein College of Medicine, 1300 Morris Park Avenue Bronx, NY 10461, USA) for providing the miRFP703 plasmid via Addgene.

---

**Supplementary Figure 1:**
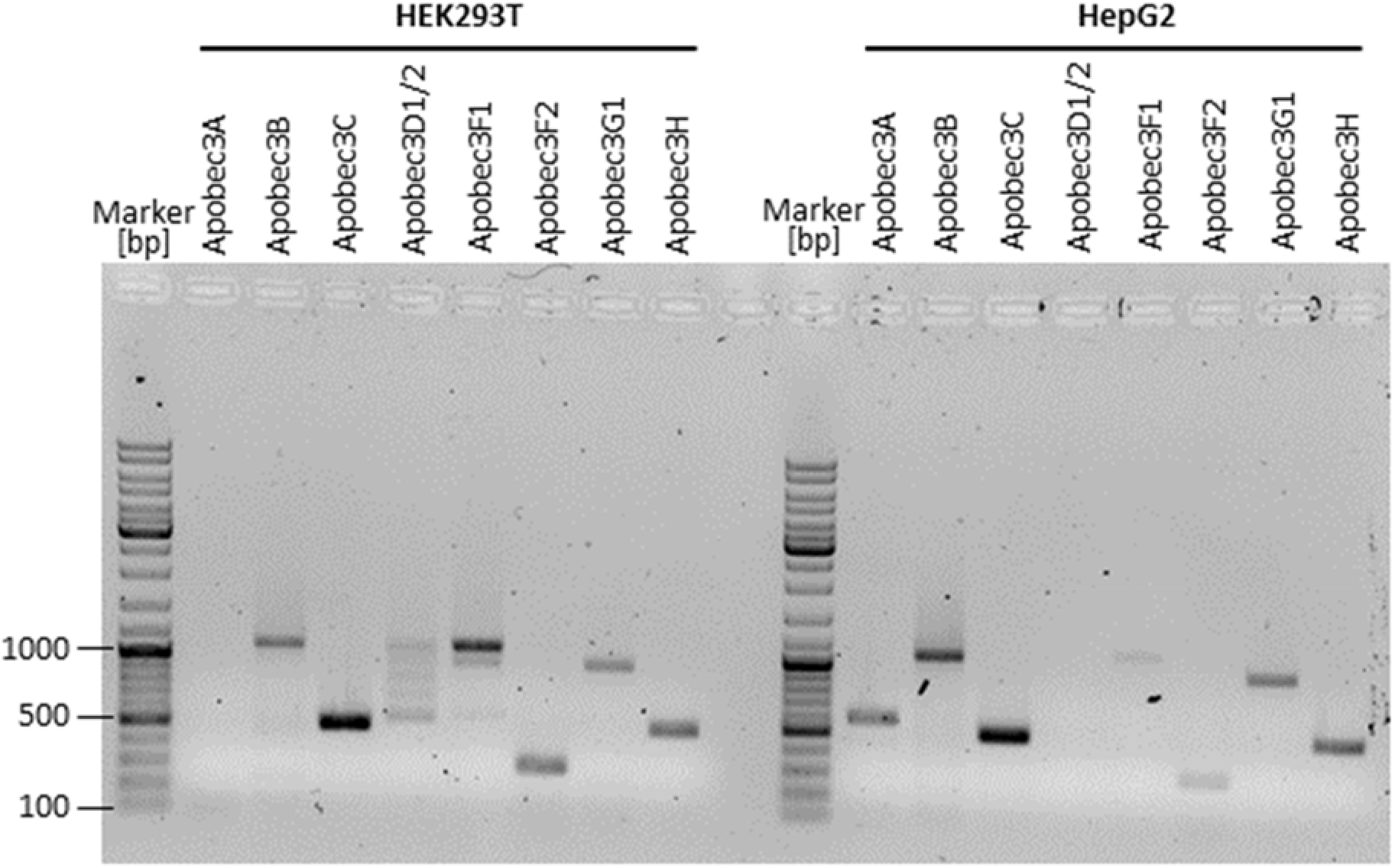
Profiling of HEK293T and HepG2 cells with regard to cell-endogenous expression of different Apobec3 variants. Same RNA-preparations as in Figure 4D and 4F were used. Oligonucleotides used for molecular profiling of these cells were as follows: Apobec3A (5’-ATCCGGGCCCAGACACTTGAT-3’ and 5’-AGTTTCCCTGATTCTGGAGAAT-3’), Apobec3B (5’-TTTGAAAACGAACCCATCCTCTAT-3’ and 5’-AGTTTCCCTGATTCTGGAGAAT-3’), Apobec3C (5’-CCTATGGGAAGCCAACGA-3’ and 5’-CTCCCGTAGCCTTCTTTTCA-3’), Apobec3D1/2 (5’-GAAAACGAACCCATCCTCT-3’ and 5’-CTCCCGTAGCCTTCTTTTCA-3’), Apobec3F1 (5’-GCCTCACTTCAGAAACACAGTGG-3’ and 5’-AGCTTGCTGTCCAGGAATAG-3’), Apobec3F2 (5’-GCCTCACTTCAGAAACACAGTGG-3’ and 5’-TGTTCCCTGCCTGCGGTCAG-3’), Apobec3G1 (5’-GCCTCACTTCAGAAACACAGTGG-3’ and 5’-AGTGAAGATGCACAGGCTCA-3’), and Apobec3H (5’-CCTCAGAAGGCCTTACTAC-3’ and 5’-CTTTATCCTCTCAAGCCGTC-3’). Please note that amplified Apobec3D1 yields a band with 1086 bp while Apobec3D2 is detectable at 534 bp.

